# PInteract: Detecting Aromatic-Involving Motifs in Proteins and Protein-Nucleic Acid Complexes

**DOI:** 10.1101/2025.07.23.666328

**Authors:** Dong Li, Fabrizio Pucci, Marianne Rooman

## Abstract

With the recent development of accurate protein structure prediction tools, virtually all protein sequences now have an experimental or a modeled structure. It has therefore become essential to develop fast algorithms capable of detecting non-covalent interactions not only within proteins but also in protein-protein, protein-DNA, protein-RNA, and protein-ligand complexes. Interactions involving aromatic compounds, particularly their *π* molecular orbitals, hold unique significance among molecular interactions due to the electron density delocalization, which is known to play a key role in processes such as protein aggregation. In this paper, we present PInteract, an algorithm that detects *π*-involving interactions in input structures based on geometric criteria, including *π*–*π*, cation–*π*, amino–*π*, His–*π*, and sulfur–*π* interactions. In addition, it is capable of detecting chains and clusters of *π* interactions as well as particular recurrent motifs at protein-DNA and protein-RNA interfaces, called *stair motifs*, consisting of a particular combination of *π*–*π* stacking, cation/amino/His–*π* and H-bond interactions. PInteract is freely available for download at https://github.com/3BioCompBio/PInteract.

## 1 Introduction

Among the various types of noncovalent interactions that stabilize the 3-dimensional (3D) structures of proteins and protein-involving complexes, interactions where one of the partners carries an aromatic ring have a status apart [1, 2]. Indeed, their London dispersion contribution to the free energy is particularly important, due to the strength of *π*-electron delocalization. In particular, interactions between two aromatic moieties, called *π − π* interactions [3], have large dispersion energies especially if the aromatic rings are parallel. When the rings are in T-shaped conformations, the electrostatic contributions are increased at the expense of dispersion contributions. Both T-shaped and stacked conformations are energetically favorable; their relative preference depends on the particular type of extracyclic atoms and on the environment. Here, we considered *π − π* interactions in which one of the partners is an aromatic amino acid (Phe, Tyr, Trp) and the other is either another aromatic amino acid or a nucleic acid base (Gua, Ade, Cyt, Thy, Ura).

In addition to *π − π* interactions, we considered cation-*π*, amino-*π*, and His-*π* interactions, where the *π*-partner is the ring of an aromatic amino acid or nucleic acid base. In cation-*π* interactions [4, 5, 6], the other partner is a positively charged amino acid Arg or Lys. In Lys-*π* interactions, the electrostatic contribution is predominant, whereas in Arg-*π* interactions the dispersion energy is also important due to the charge delocalization on the planar guanidinium group of the Arg side chain. In amino-*π* interactions [5, 7], sometimes also called amide-*π*, the second partner is the neutral side chain of Asn or Gln, which is the planar and polarizable formamide group; the electrostatic contribution is here small and the dispersion energy is the main stabilizing force. The His-*π* interaction [8] is, in essence, contained in other subtypes according to the protonation state of the His: when neutral, His-*π* can be considered as a *π*-*π* interaction, and when protonated, as a cation-*π* interaction. This makes His-*π* a special and versatile pH-dependent interaction that plays a unique role in function and molecular recognition [9, 10]. We chose to treat it as a separate form of *π* interaction.

The last *π*-involving type of interactions we considered is sulfur-*π* interaction [11, 12], also called thiol-*π*, which links an aromatic moiety of an amino acid or nucleic acid base with a sulfur atom of Met or Cys side chains. Such interactions have been shown to be stabilizing, though less than other *π* interactions [13], and to be sometimes of functional importance [14].

Different *π* interactions are often found to combine into larger motifs, such as chains or clusters of *π−π* interactions [15] and of cation/amino/His-*π* interactions [16]. For example, a positively charged ion can be sandwiched between two aromatic rings, or an aromatic ring can be sandwiched between two positively charged groups. Moreover, recurrent motifs called stair motifs, which mix *π − π*, cation-*π*, amino-*π*, His-*π* and H-bonds, are also commonly found at protein-DNA interfaces [16, 17].

There has been great success in studying the importance of aromatic-involving interactions in biological processes [18, 19, 20, 21]. For example, cation-*π* and *π*-*π* interactions have been shown to dictate the phase transition in liquid-liquid phase separation [22]. Furthermore, by mutating some residues in histones into Trp, thus increasing the number of cation-*π* interactions, researchers managed to improve the affinity between histones and their reader proteins, which further enhances downstream genetic regulations [23]. Moreover, a mutation from Phe to Thr in BanLec, a banana protein which is regarded as a potential antiviral agent and works by binding to glycan molecules via *π*-*π* stacking, was demonstrated to reduce the protein’s antiviral competence as the mutation disrupted the capacity of glycan binding of BanLec. Sulfur-*π* interactions, despite a relatively lower appearance frequency, have been pointed out for contribution to protein stability [24]. Finally, His-*π* interactions, being interchangeable among different electrostatic states, has been shown to impact protein activity according to the environmental salt concentration.

Several databases of aromatic interactions in proteins have been publicly released, including CAD [25] and A^2^ID [26], each encompassing different types of aromatic interactions. Despite these advancements, there is still a lack in publicly released computational tools allowing the identification of different types of aromatic interactions. To fill this gap, we developed PInteract, a program written in C, which is able to quickly and accurately localize aromatic-involving interactions in input structures by a criteria combining both distances and angles, which is more chemically reasonable than distance-only tools [27]. With a wide range of supported types of input structures, including protein monomers, multimers, and protein-nucleic acid complexes, we expect PInteract to be an efficient tool in the calculation and analysis of aromatic interactions in biological molecules.

## 2 Materials and Methods

### 2.1 Types of interactions

In the interactions we considered, the first partner is an aromatic moiety of an amino acid (Phe, Tyr, Trp, His) or a nucleic acid (Gua, Ade, Cyt, Thy, Ura). The second partner is an amino acid which is either aromatic (Phe, Tyr, Trp), contains a sulfur atom (Met, Cys), carries a net positive charge (Lys, Arg) or a partial one (Asn, Gln), or His that is both aromatic and sometimes positively charged. According to the type of partner, the aromatic-involving interactions are referred to as *π − π*, sulfur-*π*, cation-*π*, amino-*π*, and His-*π*, respectively.

### 2.2 Design principles

PInteract detects *π* interactions using geometric criteria derived from energetic principles established through quantum chemical studies (see, e.g., [5]). The interaction energy, defined as Δ*E* = *E_A−B_ −* (*E_A_* + *E_B_*), serves as a unified descriptor for diverse *π* interactions, where *E_A−B_* is the total energy of the interacting partners *A* and *B*, and *E_A_* and *E_B_* are the energies of the isolated components. A negative Δ*E* indicates a stabilizing interaction. Key factors that determine the geometry and strength of *π* interactions include:

- The distance between the two interacting partners. Sometimes, this distance is defined as the distance between the centroids of the two interacting functional groups. In other studies, like here, it is defined as the closest distance between any two atoms of the two functional groups. We chose this distance definition because the considered aromatic moieties carry various substituents which influence how the ring interacts with other partners and breaks the symmetry around the ring’s center. Similarly, the electron delocalization in other functional groups is better captured by selecting the closest atom rather than the functional group’s centroid.
- The *α* angle; it is typically defined as the angle between the vector linking the centroids of both partners and the vector normal to the aromatic ring of the aromatic partner or of one of the aromatic partners in the case of *π − π* interactions. The complement of *α*, i.e. 90*^◦^ − α*, is called elevation angle and is sometimes also used.
- The *β* angle; it only applies to interactions in which the two functional groups are planar, i.e. contain an aromatic moiety (Phe, Tyr, Trp, His), a guanidinium group (Arg) or an formamide group (Asn, Gln). It is defined as the angle between the vectors normal to the planes of the two interacting functional groups and measures the degree of parallellism between the planes.

The energy values of the interacting partners vary significantly according to their interaction geometries. Optimal Δ*E* values are observed for inter-partner distances of 2.5-4.0 Å for cation-*π* and amino-*π* interactions [5, 28, 29, 25, 18], about 6.0 Å for sulfur-*π* [11, 30], and up to about 5.0 Å for *π*-*π* interactions [2]. In addition to the distance, the angular parameter *α* constraints the interaction geometry and modulates the interaction energies. It allows the specification that the functional group of one partner is positioned above (or below) the aromatic moiety of the other, thereby enabling interaction with its *π* orbital. The *β* angle further influences the interaction energy by defining whether the interacting molecular planes adopt a stacked or T-shaped geometry.

### 2.3 Technical implementation

#### 2.3.1 Functional groups

The *π* interactions between functional groups are defined using geometric criteria. The atoms considered to be part of the functional groups are the following:

- Aromatic moiety: all atoms that make up the aromatic ring or rings;
- Positive charge: NH1, NH2, CZ and NE for Arg; and NZ, CE, H1 for Lys; ND1, CE1, NE2 for His;
- Partial positive charge: NE2, CD and OE1 for Gln; ND2, CG and OD1 for Asn;
- Sulfur: SG for Cys; SD for Met.

By convention, partner 1 is taken as the aromatic moiety, or one of the aromatic moieties in the case of *π − π* interactions.

#### 2.3.2 Distance criterion

The first criterion is a distance threshold, *d*_max_, which specifies the maximum allowed separation between atoms involved in the interaction. Specifically, we compute the smallest distance *d* between any atom of the functional group of partner 1 and any atom of the functional group of partner 2, specified in Section 2.3.1. This is illustrated in Figure 1a. For the interaction to be considered valid, this distance must satisfy *d ≤ d*_max_ where:

- *d*_max_ = 5.0 Å for *π*-*π* interactions;
- *d*_max_ = 6.0 Å for sulfur-*π* interactions;
- *d*_max_ = 4.5 Å for cation-*π*, amino-*π*, and His-*π* interactions.

**Figure 1:**
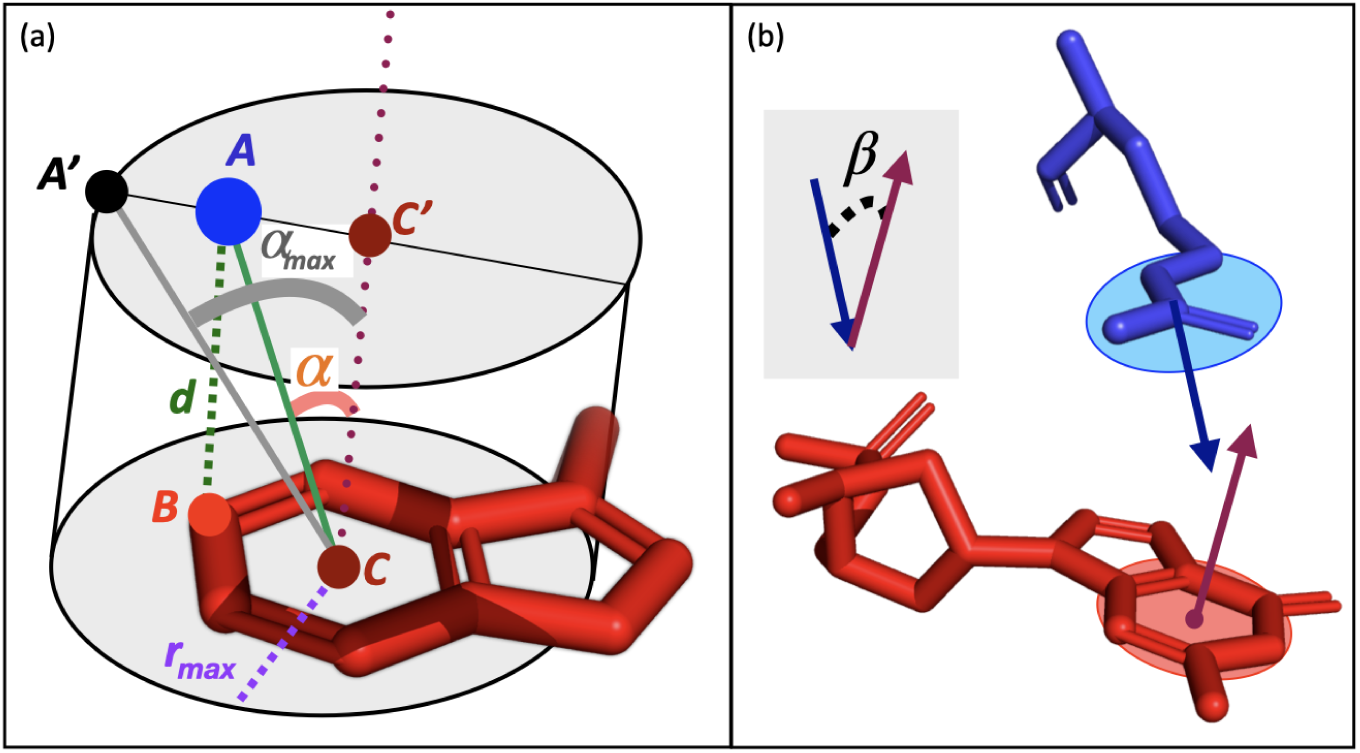
Geometric definition of *π*-involving interactions. (a) The aromatic moiety of partner 1 (here Trp) is in red; the centroid of its 6-atom ring is labeled *C*. The closest atoms between the two partners are atom *B* of the 6-atom aromatic ring and atom *A* of the functional group of partner 2, represented by a blue ball; their distance is equal to *d*. *A* must be inside a cylinder, whose lower base is a circle of radius *r_max_*centered on *C* and its upper base passes through *A* and is centered on *C^′^*. The angle between vector 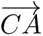 and vector 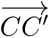 which is normal to the cylinder’s bases, is called *α*. For *A* being inside the cylinder, *α* must be *≤ α_max_*, where *α_max_* is the angle between the normal vector 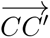 and vector 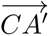, where *A^′^* is on the upper base’s edge and on the diameter passing through *A* and *C^′^*. (b) Partner 1 (Trp) is in red and partner 2 (Arg), in blue. The planes formed by the 6-atom aromatic ring of Trp and by Arg’s guanidinium group are shown in red and blue, respectively, as well as the vectors normal to these planes. The angle between these vectors, called *β*, taken between 0*^◦^* and 90*^◦^*, is shown on a gray background and measures the deviation from the parallelism between the two planes.

These distance thresholds are provided by default, but they can easily be modified by the user. This is particularly important when using low-resolution or modeled protein 3D structures, in which the positioning of the side chain atoms is not sufficiently accurate.

If an aromatic partner consists of a fused double-ring system, one ring with 5 atoms and the other with 6, as is the case for Trp, Ade and Gua, we determine whether the closest distance involves an atom that is closer to the centroid of the 6-atom ring or of the 5-atom ring. In the former case, we focus on the 6-atom ring for checking whether the other geometrical criteria are satisfied and in the latter case, on the 5-atom ring.

#### 2.3.3 Angle criterion

To ensure that the interacting partner is positioned within the effective electronic range of the *π* orbital, and thus that the interaction is *π*-mediated, we apply an angular constraint based on a cylindrical model, illustrated in Figure 1a. The cylinder’s base is centered on the geometric center *C* of the aromatic ring of partner 1 (or of the ring closest to partner 2 in the case of a double aromatic moiety), and its radius is equal to *r_max_* taken to be :

- *r_max_* = 2 *r* for cation-*π*, His-*π*, amino-*π* and sulfur-*π* interactions;
- *r_max_* = 3 *r* for *π*-*π* interactions.

where *r* is the radius of the aromatic ring. The height of the cylinder is such that the closest atom of the functional group of partner 2 (noted *A*) is included in the cylinder’s second (”upper”) base. If this is impossible, thus if *A* remains outside the cylinder regardless of its height, the interaction is not considered as a *π* interaction.

The *r_max_* threshold can be translated into an angle criterion. Indeed, for *A* to be in the cylinder, the angle *α* formed by vector 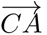, linking the aromatic moiety’s center to *A*, and the vector normal to the aromatic ring (taken in the direction of *A* so that *α ≤* 90*^◦^*) must have a value that is lower than a maximum value, called *α_max_*. This value is reached when *A* is on the cylinder’s edge, as shown in Figure 1a. Note that the *α_max_* threshold value depends on the distance *d* between the the 2 partners, unlike the *r_max_* threshold.

The *r_max_* thresholds are provided by default in the PInteract program but, like for the distance thresholds, they can easily be changed by the user. Since the asymmetrically distributed electron cloud can extend beyond the aromatic ring’s atomic boundaries, using a larger radius ensures inclusion of geometries where the functional group of partner 2 is laterally offset from the ring’s centroid of partner 1 but still engages in energetically significant *π* interactions.

#### 2.3.4 Plane parallelism angle

In the case the functional groups of the two partners are planar, as in Arg-*π*, amino-*π*, His-*π*, and *π*-*π* interactions (see Section 2.2), we compute the smallest angle *β* between the vectors that are normal to the planes of the interacting functional groups, as illustrated in Figure 1b for an Arg-Trp interaction. The *β* value is equal to 0*^◦^* when the two planes are parallel, or equivalently when the two partners are in stacked conformation, and equal to 90*^◦^* when the planes are perpendicular or in T-shaped conformation.

### 2.4 Structure datasets’ construction

To apply PInteract at proteome scale, we set up several non-redundant and good-resolution structure datasets. The *D*_monomer_ dataset was collected from X-ray structures of protein monomers using the PISCES server [31]. We used the following filters: no missing main-chain atomic coordinates, resolution *≤* 2.5 Å, R-factor *≤* 0.25, sequence length between 40 and 10,000 residues, and pairwise sequence identity *≤* 30%. *D*_monomer_ contains a total of 5,590 proteins.

We also set up three datasets of protein-protein complexes. The first, denoted *D*_dimer_, contains heterodimers; the second one, *D*_TCRpMHC_, contains complexes between a T-cell receptor (TCR) and a peptide-bound major histocompatibility complex (pMHC); and the third one, *D*_AbAg_, contains complexes between an antibody and an antigen. Complexes from *D*_dimer_ were collected and curated using the PISCES server. The TCRpMHC complexes of *D*_TCRpMHC_ were obtained from the TCR3d database [32, 33], and the antibody-antigen complexes of *D*_AbAg_, from the SAbDab database [34, 35, 36], using the same filters as for *D*_monomer_. These three datasets contain 1,037 heterodimer structures, 51 TCR-pMHC structures and 495 antibody-antigen structures, respectively.

To compute the frequency of occurrence of *π* interactions in these datasets, we divided the number of occurrences of a given type by the total number of interactions. More specifically, for *D*_monomer_, the total number of interactions was estimated as the number of residue pairs in contact, defined as having at least one heavy atom from each residue’s side chain within a maximum distance of 5 Å. For *D*_dimer_, the frequencies were obtained in the same way except that only contacts across the protein-protein interface were considered. In *D*_AbAg_, where the complexes typically consist of three chains, the considered interactions link the antigen and antibody chains. In *D*_TCRpMHC_, where the complexes consist of four or five chains, the interactions considered are between the TCR chains and the pMHC molecule.

## 3 Results

### 3.1 The PInteract algorithm

We developed the PInteract algorithm to identify *π*-involving interactions in proteins, protein-protein complexes, and interfaces between proteins and DNA, RNA, or nucleic acid base-containing ligands. These interactions include *π − π*, sulfur-*π*, cation-*π*, amino-*π*, and His-*π* interactions, which are determined based on the geometric criteria detailed in Section 2.3.

The input to PInteract is the name of a directory that contains structure files in PDB format [37]. The output consists of three files: PInteract.csv, a table of individual *π* interactions; PInteract1.csv, a table of *π*-chains and stair motifs; and PInteract.txt, a human-readable text file containing all the content from PInteract.csv and PInteract1.csv, with comments and explanations prefixed by # to aid understanding. PInteract is implemented in C and is extremely fast, enabling the detection of *π* interactions at the proteome scale. The code can be freely downloaded from our GitHub repository: https://github.com/3BioCompBio/PInteract. All technical details are provided in this repository.

### 3.2 Examples of π-involving interactions

Here, we provide examples of PInteract applications to identify specific geometric arrangements of *π*-involving interactions.

The first example of PInteract’s application is illustrated in Figure 2a. A cation-*π* and a *π − π* interaction were found at the interface between hen egg lysozyme and the heavy chain of an antibody. These two interactions are generally particularly important in antibody-antigen recognition [38]. In this specific case, for example, their disruption has been experimentally shown to reduce the antibody-antigen binding affinity by at least two orders of magnitude [39].

**Figure 2:**
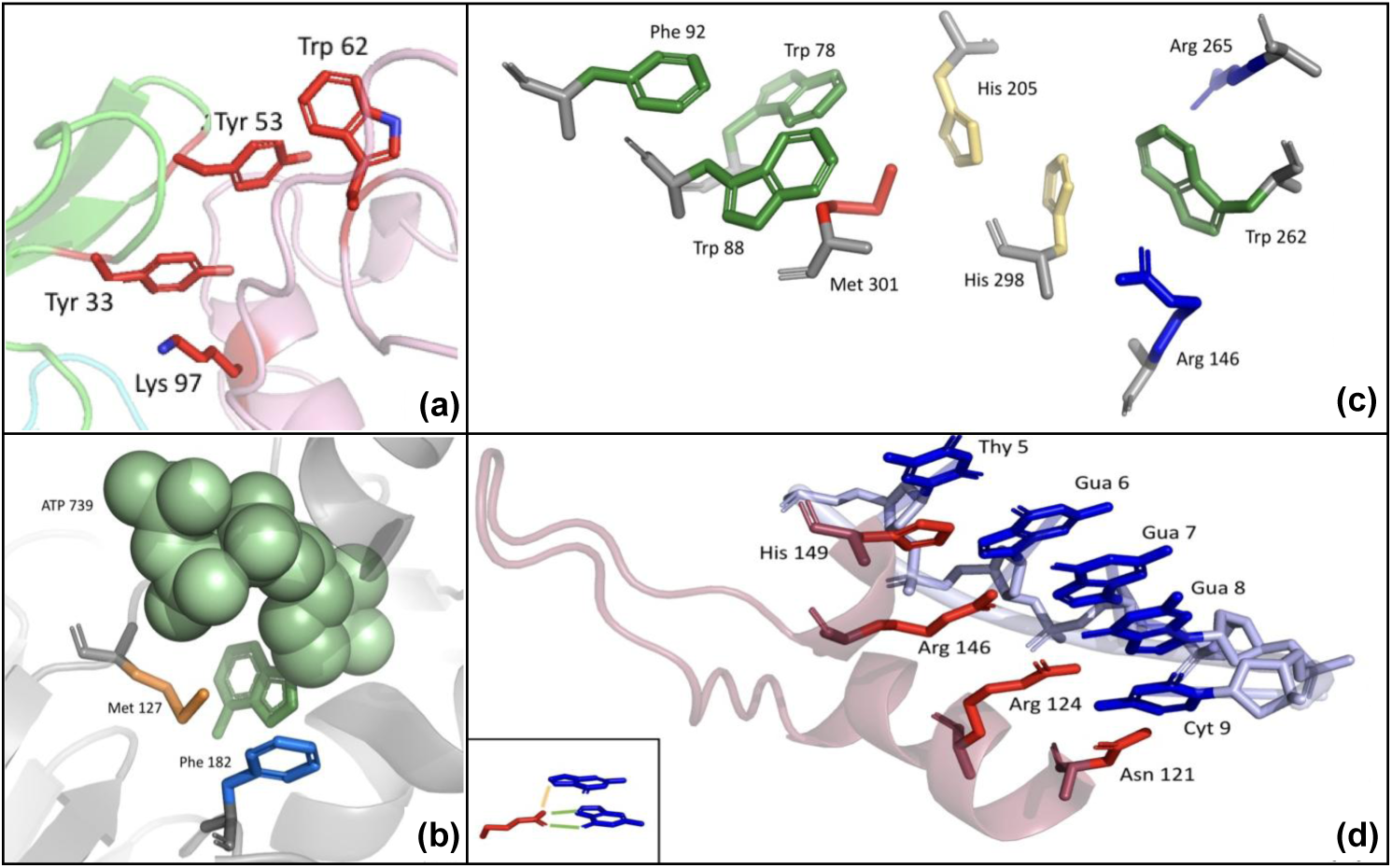
Illustration of *π*-involving interactions. (a) Example of *π* interactions at a protein-protein interface. Cation-*π* interaction between Lys Y97 and Tyr H33 and *π*-*π* interaction between Tyr H53 and Trp Y62 at the interface between a hen egg lysozyme and its cognate antibody (PDB ID: 2QDJ). Chain Y is the antigen depicted in pink; chain H is the antibody’s heavy chain and is shown in green; the light chain is shown in cyan. (b) Example of protein-ligand *π* interactions. ATP molecule C739 (in green spheres with its Ade nucleobase in green sticks), Phe B182 (in blue sticks) and Met B127 (in orange sticks) in an NAD kinase from *Archaeoglobus fulgidus* (PDB ID: 1Z0S). (c) Example of *π*-chains. Residues Trp 78, Trp 88, Phe 92, Arg 146, His 205, Trp 262, Arg 265, His 298, Met 301 in chain A of a proteobacterium’s carboxylesterase (PDB ID: 1QLW). Aromatic side chains are in green, sulfur-containing side chains in red, histidines in yellow, and positively charged side chains in blue. Backbone atoms are in gray. (d) Example of stair motifs at the protein-DNA interface. Nucleic acid bases Thy B5, Gua B6, Gua B7, Gua B8, Cyt B9 and residues His A149, Arg A146, Arg A124 and Asn A 121 in a zinc finger-DNA complex in *Mus musculus* (PDB ID: 1A1G). Nucleic acid bases are depicted as blue sticks and amino acid side chains, as red sticks. In the lower left corner, the H-bonds are depicted as green lines and the cation-*π* interaction as an orange line.

PInteract is also able to detect *π* interactions with ligands that contain a nucleic acid base, such as ATP or GPT. An example of such a protein-ligand *π* interactions is shown in Figure 2b. A Met and a Phe residue of a NAD kinase make a sulfur-*π* and a *π − π* interaction, respectively, with the Ade base included in the ATP molecule.

Successive *π*-involving interactions identified by our algorithm are referred to as *π*-chains in the output files. These chains represent continuous networks of interacting *π*-systems, which may play a structural or functional role in stabilizing molecular assemblies. They can be successive interactions of the same type, e.g., clusters of *π*-*π* interactions or a double cation-*π* interaction where the cation is sandwiched between aromatic rings or where an aromatic moiety is sandwiched between two cations. They can also mix different types of *π* interactions, e.g. a *π − π* interaction followed by a cation-*π* interaction. An example is given in Figure 2c, which shows a *π*-chain formed of three *π − π*, two cation-*π* and three sulfur-*π* interactions in a bacterial esterase from an *Alcaligenes* species.

Another type of mixed interactions identified by PInteract is the stair motif [16, 17], recurrently observed at protein-DNA or protein-RNA interfaces. These motifs consist of two stacked nucleic acid bases making a *π − π* interaction, and an amino acid that forms both a cation-*π*, amino-*π* or His-*π* interaction with one of the nucleic acid bases and an H-bond with the next base along the base stack. For PInteract to be able to detect such motifs, the HBPLUS algorithm [40] that detects H-bonds must be run before. An example of four successive stair motif is shown in Figure 2d, where five stacked nucleobases make H-bonds and His-*π*, cation-*π* and amino-*π* interactions with a His, two Arg and an Asn of a zinc finger protein.

### 3.3 Large-scale analysis of π-involving interactions

In order to show the applicability of our tool for large-scale analysis of *π*-involving interactions, we investigated the relative frequency of each type of *π* interaction across the proteins in the four *D* datasets described in Section 2.4: monomeric proteins, heterodimers, antibody-antigen complexes, and TCR-pMHC complexes. These frequencies were computed as the number of interactions identified by PInteract divided by the total number of interactions in the protein (see Section 2.4 for details).

As shown in Table 1 and Figure 3, antibody-antigen interfaces contain the largest proportion of *π* interactions on the average, followed by TCR-pMHC interfaces; heterodimer interfaces and monomers have lower, almost identical, proportions. More specifically, antibody-antigen interfaces contain by far the largest proportion of cation-*π* and *π*-*π* interactions. The role of these interactions in antibody-antigen binding is well known [41].

**Figure 3:**
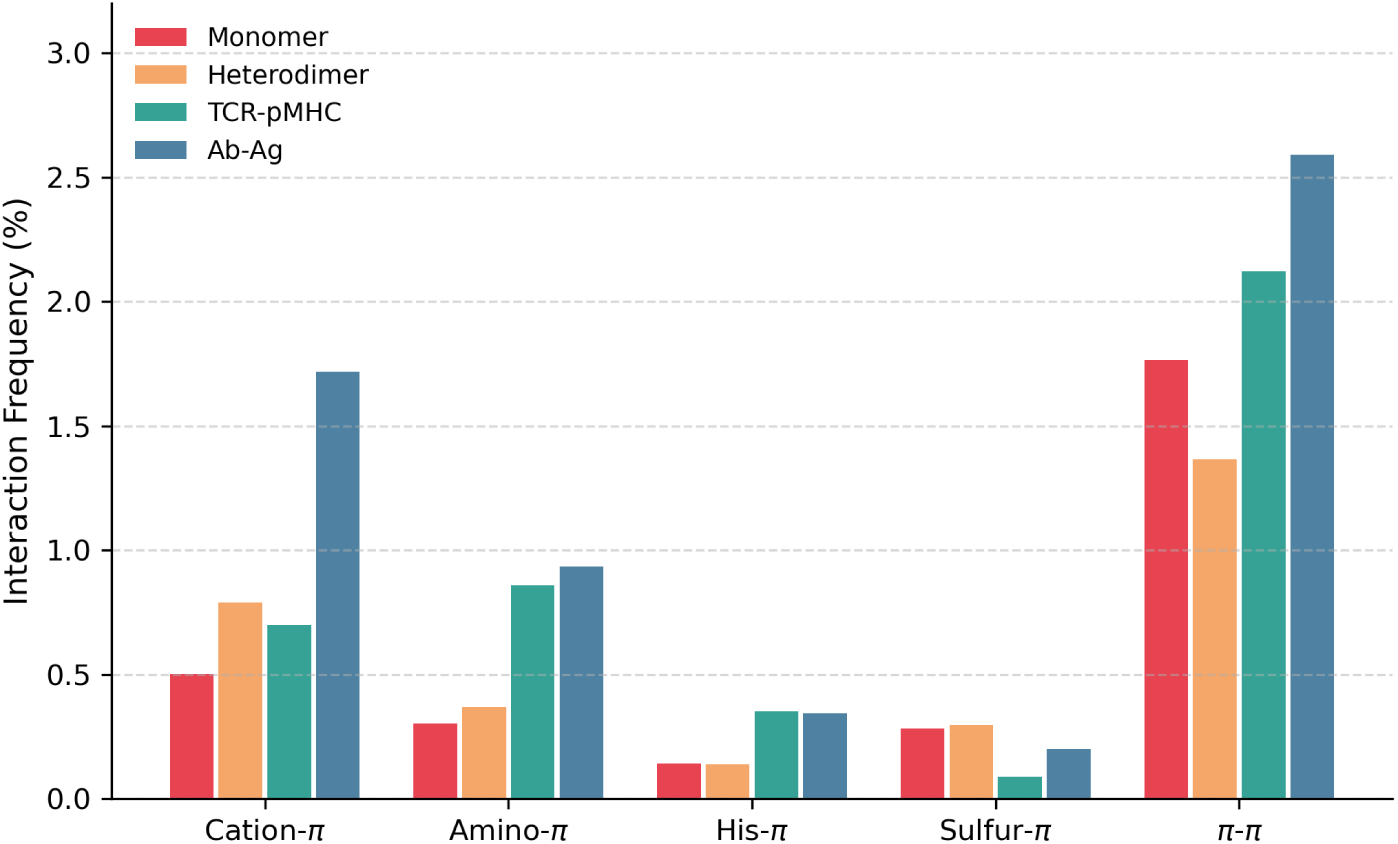
Comparison between the average relative frequency of each type of *π* interactions in the datasets *D*_monomer_, *D*_dimer_, *D*_TCRpMHC_, and *D*_AbAg_. The relative frequencies were computed as the number of interactions identified by PInteract divided by the total number of interactions in the protein, as defined in Section 2.4. For the heterodimers, only the interactions occurring at the interface were counted; for antibody-antigen complexes, only interactions that link one of the two antibody chains with the antigen; and for TCR-pMHC, only interactions that link one of the two TCR chains with the pMHC molecule.

**Table 1:**
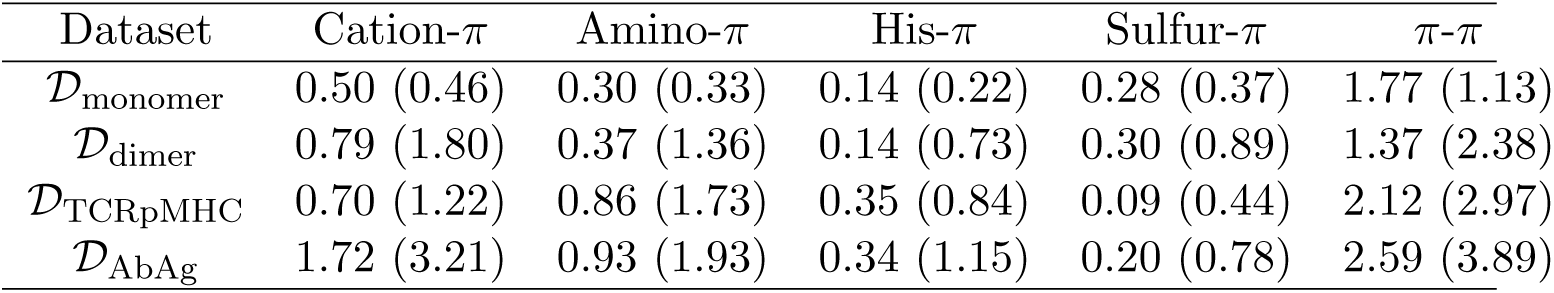
Average relative frequencies of *π* interactions in the set of monomers *D*_monomer_ and in the sets of protein-protein complexes *D*_dimer_, *D*_TCRpMHC_, and *D*_AbAg_, computed as the number of interactions identified by PInteract divided by the total number of interactions in the protein (see Section 2.4). The standard deviation is given in parentheses. The associated p-values are given in Supplementary Table S1.

| Dataset | Cation-π | Amino-π | His-π | Sulfur-π | π-π |
| --- | --- | --- | --- | --- | --- |
| $\mathcal{D}_{\text{monomer}}$ | 0.50 (0.46) | 0.30 (0.33) | 0.14 (0.22) | 0.28 (0.37) | 1.77 (1.13) |
| $\mathcal{D}_{\text{dimer}}$ | 0.79 (1.80) | 0.37 (1.36) | 0.14 (0.73) | 0.30 (0.89) | 1.37 (2.38) |
| $\mathcal{D}_{\text{TCRpMHC}}$ | 0.70 (1.22) | 0.86 (1.73) | 0.35 (0.84) | 0.09 (0.44) | 2.12 (2.97) |
| $\mathcal{D}_{\text{AbAg}}$ | 1.72 (3.21) | 0.93 (1.93) | 0.34 (1.15) | 0.20 (0.78) | 2.59 (3.89) |

The TCR-pMHC binding interfaces also contain a large proportion of *π*-*π* interactions, intermediate between antibody-antigen and other heterodimer interfaces, whereas the proportion of cation-*π* interactions is the same as in other heterodimer interfaces. The proportion of amino-*π* and His-*π* is identical in TCR-pMHC and antibody-antigen interfaces and larger than in heterodimer interfaces and in monomers. In contrast, sulfur-*π* interactions are rare in general, and even rarer in TCR-pMHC and antibody-antigen interfaces. These tendencies are statistically significant as shown in Supplementary Table S1. Note, however, that the standard deviation is very high (Table 1) and thus that the variability between proteins is important.

To investigate the characteristic distances between the functional groups involved in *π* interactions and evaluate the relevance of our predefined distance thresholds *d_max_*, we analyzed the distribution of distances *d* (defined in Section 2.3.2) for cation-*π*, amino-*π*, His-*π*, sulfur-*π*, and *π*-*π* interactions detected in dataset *D*_monomer_. The results are shown in Figure 4. Clearly, as a general trend, the interaction distances consistently exceed 3 Å. The *d*-distributions exhibit characteristic peaks at different values according to the interaction types: 3.6 Å for cation-*π*, amino-*π*, and His-*π* interactions, and 3.8 Å for sulfur-*π* and *π*-*π* interactions. Interestingly, the peak is less marked for sulfur-*π* interactions, showing that the geometry of this type of interaction is less constrained. Our analysis also confirms that the chosen thresholds for each interaction type are reasonable, as these are located near the tail of the distribution, corresponding to low-density regions. Note again that the different distance thresholds can be modified by the users.

**Figure 4:**
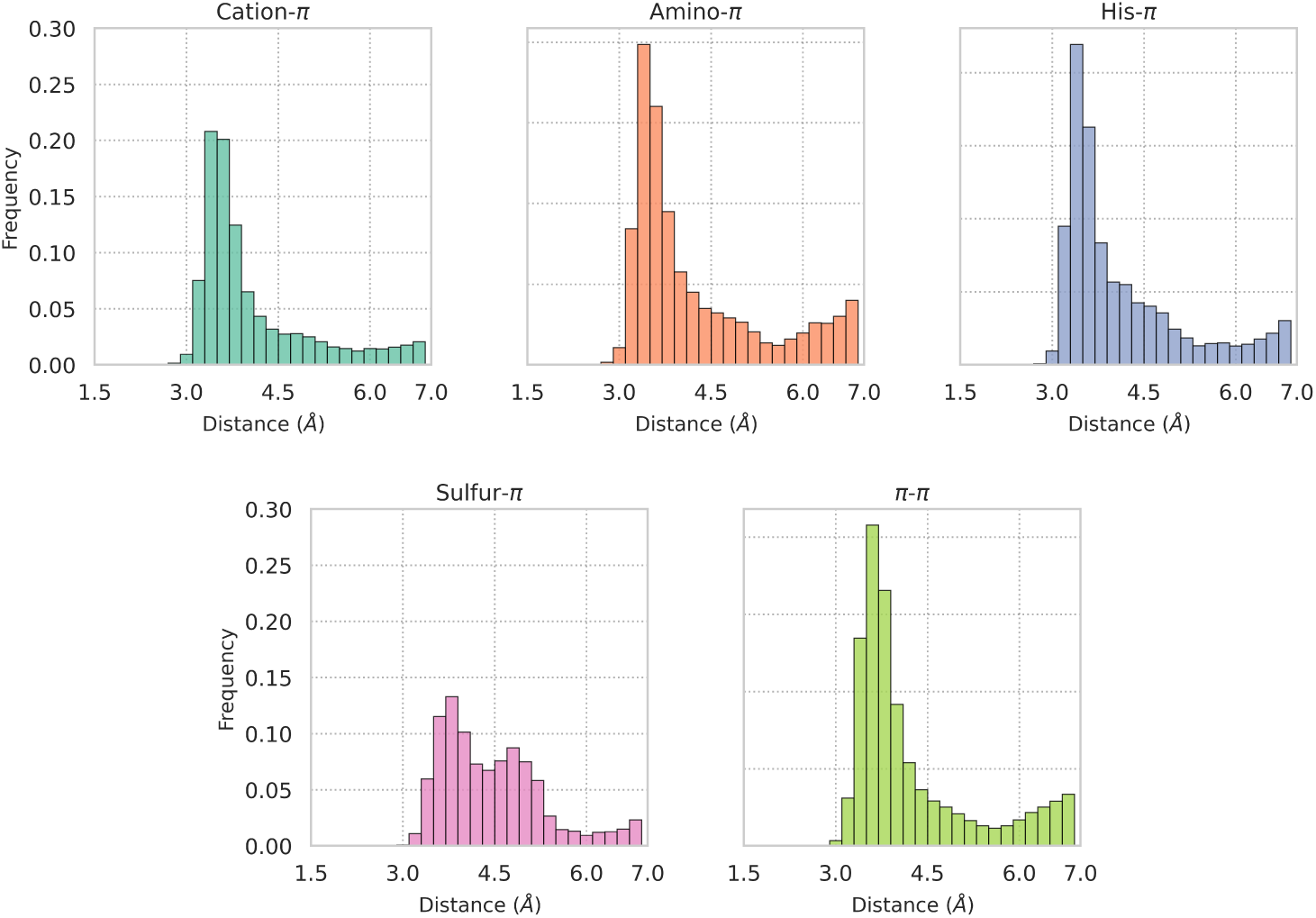
Distribution of the distances *d* between functional groups of *π*-involving interactions, defined in Section 2.3.2, for all considered types of *π* interactions detected in the *D*_monomer_ dataset.

We also analyzed the distribution of *α* angle values, defined in Section 2.3.3, which represent the position of the functional group of partner 2 above (or below) the aromatic ring of partner 1. A value of *α* = 0 indicates that this functional group is just above (or below) the aromatic ring’s center. We plotted the frequency of (1 *−* cos *α*) values, which represent the lateral displacement relative to the ring’s center, for the different interaction types observed in *D*_monomer_. As shown in Figure 5, all (1 *−* cos *α*) values have an almost equal frequency up to *α* angles of about 30° (1 *−* cos 30*^◦^ ≈* 0.13) for all interaction types except *π − π*. This indicates that the formation of *π* interactions requires the interacting partner to be clearly above (or below) the aromatic plane, where the *π*-orbital is situated; however, no clear preference for a particular displacement from the ring’s center is observed, which reflects a certain plasticity of the interaction geometry. For *α* angles larger than about 45° (1 *−* cos 45*^◦^ ≈* 0.23), the frequency of all non-*π*-*π* interactions totally vanishes. The fact that the decrease is gradual is related to the fact that the angle threshold value *α_max_*depends on the distance *d* between interacting partners and is thus not clear-cut.

**Figure 5:**
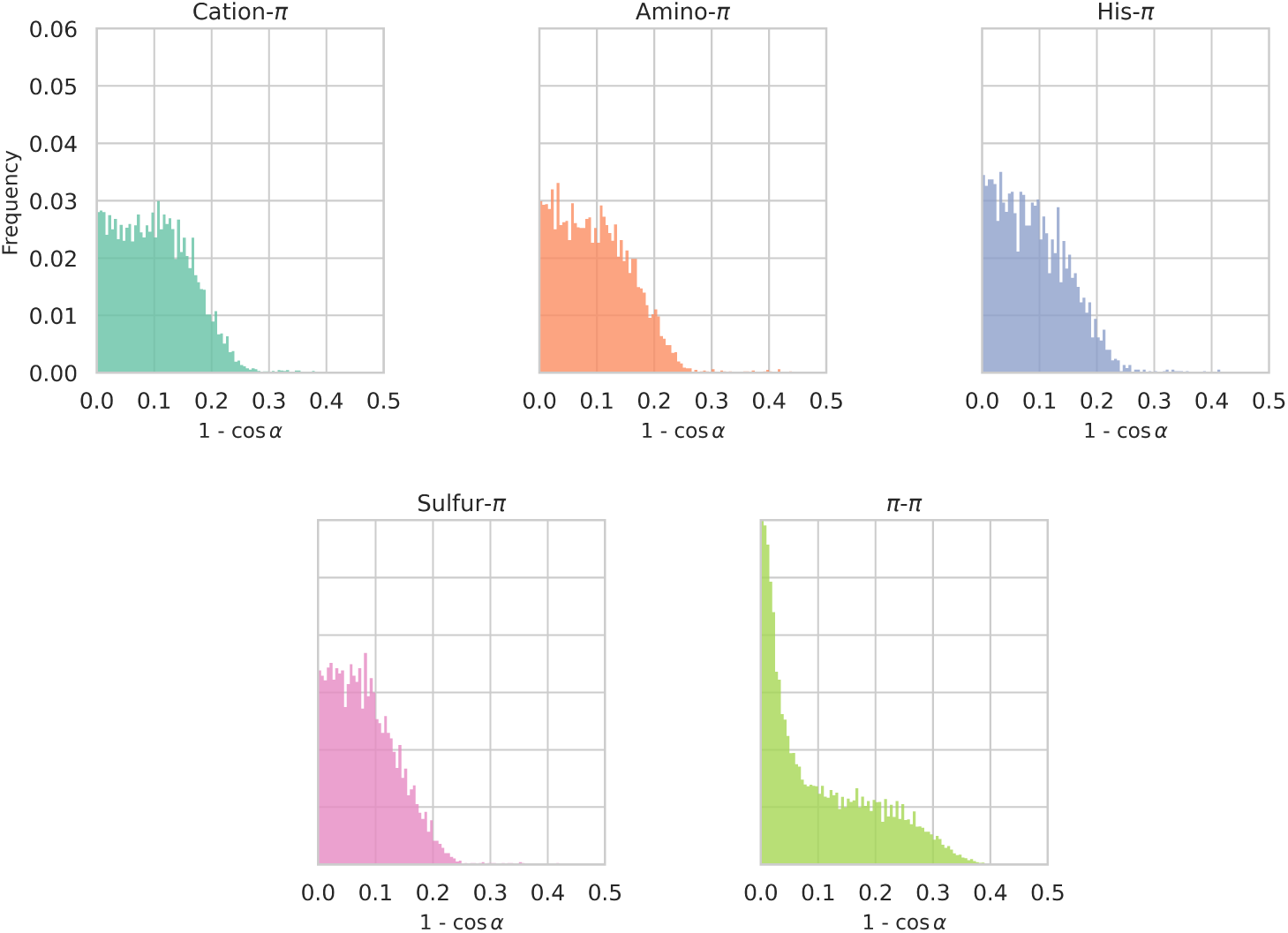
Distribution of the values of the lateral displacement with respect to the aromatic rings’ center, (1 *−* cos *α*), in which the *α* angle measures the location of the functional group of interacting partner 2 above (or below) the plane of the aromatic ring of partner 1 (see Section 2.3.3), for different types of *π* interactions detected in the *D*_monomer_ dataset. The smaller the value of (1 *−* cos *α*), the more directly the functional group of partner 2 is positioned above (or below) the center of the aromatic ring. See Supplementary Figure S1 for the corresponding *α* angle distributions.

In contrast, *π*-*π* interactions exhibit a clear preference for small displacement values, with *α* values between 0*^◦^* and about 18*^◦^* (0 *≤* 1*−*cos 18*^◦^ ≤* 0.05). This means that the closest atom of the aromatic ring of partner 2 is preferably situated above the centroid of the aromatic ring of partner 1. It has however to be noted that these displacements are not computed between the centroids of the two interacting aromatic rings, but between an atom of one aromatic ring and the center of the other ring. This can explain the over-representation of small *α* angles. Also, larger displacements are observed for this type of *π*-interactions, up to (1 *−* cos 53*^◦^ ≈* 0.40); this is due to the larger *α_max_*value accepted for this type of interaction (Section 2.3.3).

Finally, we computed the distribution of *β* angles, or rather of the (1 *−* cos *β*) displacement values, where *β* is the angle between the planes of the functional groups of the two interacting partners, when these groups are planar (Section 2.3.4). As shown in Figure 6, this distribution varies markedly depending on the type of *π* interaction. For Arg-involving cation-*π* interactions, smaller (1*−*cos *β*) values — corresponding to parallel, stacked conformations — are strongly preferred. A similar but slightly less marked trend is observed for amino-*π* and His-*π* interactions. To an even much lesser extent, this trend is also observed for *π*-*π* interactions. However, these interactions tend to accommodate a wide range of conformations, from stacked to T-shaped, with almost no preference, suggesting a high plasticity in interaction geometry. Note that these trends are similarly observed in the *β*-angle distributions of antibody-antigen complexes, TCR-pMHC, and other heterodimer complexes.

**Figure 6:**
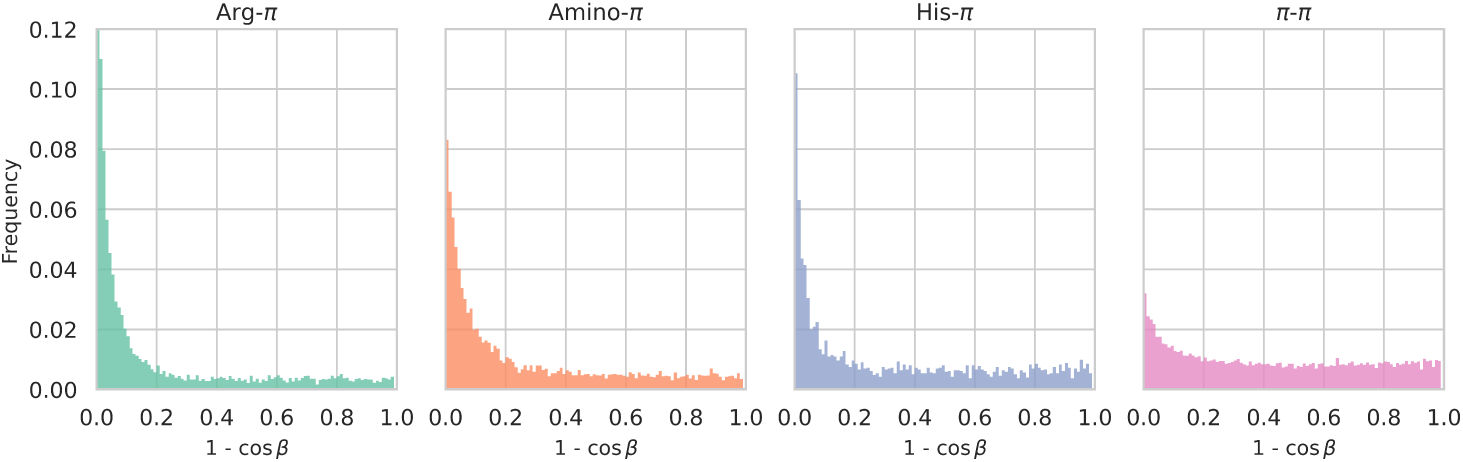
Distribution of 1 *−* cos *β* values, in which the *β* angle measures the degree of parallelism between the interacting planar functional groups (see Section 2.3.4), for different types of *π* interactions detected in the *D*_monomer_ dataset. Parallel or stacked conformations correspond to *β* = 0° and 1 *−* cos *β* = 0; perpendicular or T-shaped conformations, to *β* = 90° and 1 *−* cos *β* = 1. See Supplementary Figure S2 for the corresponding *β* angle distributions.

To further evaluate whether these observed preferences of *α* and *β* angles are preserved in computationally predicted structures, we applied PInteract to selected subsets of the AlphaFold Protein Structure Database (AlphaFoldDB) [42], as described in Supplementary Section S3. As shown in Supplementary Figures S3 and S4, the resulting distributions of *α* and *β* angles have a high degree of consistency with those observed in experimentally solved crystal structures, confirming that models predicted by AlphaFold [43] largely retain the geometric characteristics of natural *π* interactions. A minor exception is the attenuated peak of the *β* angle distribution for small *β* values, suggesting a mild underestimation by AlphaFold of the preference for stacked conformations.

### 3.4 Relationship between π interactions and protein solubility and aggregation

Solubility and aggregation are important protein properties that have huge impacts on protein fitness. Molecular mechanisms underlying protein solubility and aggregation have attracted a lot of attention but are still challenging to explain. They are complex properties that arise from the interplay of various forces in both the folded and unfolded or aggregated states. One of the results in this context is that interactions known to decrease solubility and favor aggregation involve delocalized *π*-electrons [44, 45, 46, 47].

Here, we investigated the role of intrachain *π*-involving interactions in protein solubility. For this purpose, we used the dataset of proteins with known structure and solubility values which was set up in [48]. This set contains curated entries from high-throughput experiments on the *E. coli* proteome [49] and *S. cerevisae* [50] using a cell-free expression system. Each entry in this dataset is a single-chain protein. We refer to this dataset as *D*_solubility_. We first computed the Pearson correlation between solubility values and the frequency of residues capable of forming *π* interactions; the results are shown in Table 3.4. The correlation is statistically significant and most strongly negative for aromatic residues (*ρ* = *−*0.17), followed by Arg (*ρ* = *−*0.09). His, Met/Cys, and Asn/Gln do not exhibit any significant solubility preference. In contrast, Lys showed a positive correlation with solubility (*ρ* = 0.16), consistent with the well-known Arg/Lys trend: soluble proteins tend to be depleted in arginine and enriched in lysine [51].

We further assessed whether the observed correlations differ when considering only residues engaged in *π* interactions, as identified by PInteract, compared to those that are not. As reported in Table 3.4, Arg, Met/Cys and aromatic residues involved in intrachain Arg-*π*, sulfur-*π*, and *π*-*π* interactions, respectively, tend to reduce solubility relative to the same residue types not participating in such interactions. Conversely, Lys exhibits the opposite trend, favoring solubility when it is not engaged in a *π* interaction. Our results thus indicate that interactions involving delocalized electronic *π*-systems appear to play a significant role in solubility mechanisms. This is further supported by the negative correlation of approximately −0.17 observed between protein solubility and the frequency of all detected *π* interactions.

**Table 2:** Pearson correlation coefficients *ρ* between protein solubility and (1) the residue frequency *F*; (2) the frequency *F_π_* of residues involved in a *π* interaction; (3) the frequency *F_non−π_* of residues not involved in a *π* interaction, computed from the *D*_solubility_ dataset. A cross (*×*) indicates that the result is not statistically significant (p-value ¿ 0.01).

|  | **Arg** | **Lys** | **Phe/Tyr/Trp** | **Asn/Gln** | **His** | **Met/Cys** |
| --- | --- | --- | --- | --- | --- | --- |
| $F$ | -0.09 | 0.14 | -0.17 | -0.09<sup>×</sup> | -0.04<sup>×</sup> | 0.00<sup>×</sup> |
| $F_\pi$ | -0.14 | -0.04<sup>×</sup> | -0.23 | -0.05<sup>×</sup> | -0.04<sup>×</sup> | -0.15 |
| $F_{\text{non-}\pi}$ | -0.07<sup>×</sup> | 0.15 | 0.00<sup>×</sup> | -0.09<sup>×</sup> | -0.02<sup>×</sup> | 0.04<sup>×</sup> |

As a further application of PInteract, we investigated *π*-involving interactions in the stabilization of fibril structures, which is still a matter of debate. Although aromatic residues are not strictly required for amyloid formation, they are known to facilitate and promote the fibril assembly process [52]. Specifically, we analyzed the set of fibril structures provided in [53]; we refer to this set as *D*_fibril_. We observed that the system is dominated by *π*-*π* interactions, which account for as much as 91% of all detected *π* interaction types. We examined the geometric characteristics of these interactions by comparing the distributions of the *α* and *β* angles in both fibrilar and monomeric states. Although no significant difference was observed in the *α* angle distributions, the *β* angle distributions differ markedly between the two sets, as shown in Figure 7. In particular, fibrilar structures exhibit an almost exclusive preference for stacked conformations, likely driven by the geometric symmetry and packing constraints of extended amyloid aggregates, as illustrated for the A*β*40 fibril in Supplementary Figure S5. In contrast, structures of soluble monomeric proteins display a much broader distribution of *β* angles (see Figure 6), reflecting a greater conformational variability.

**Figure 7:**
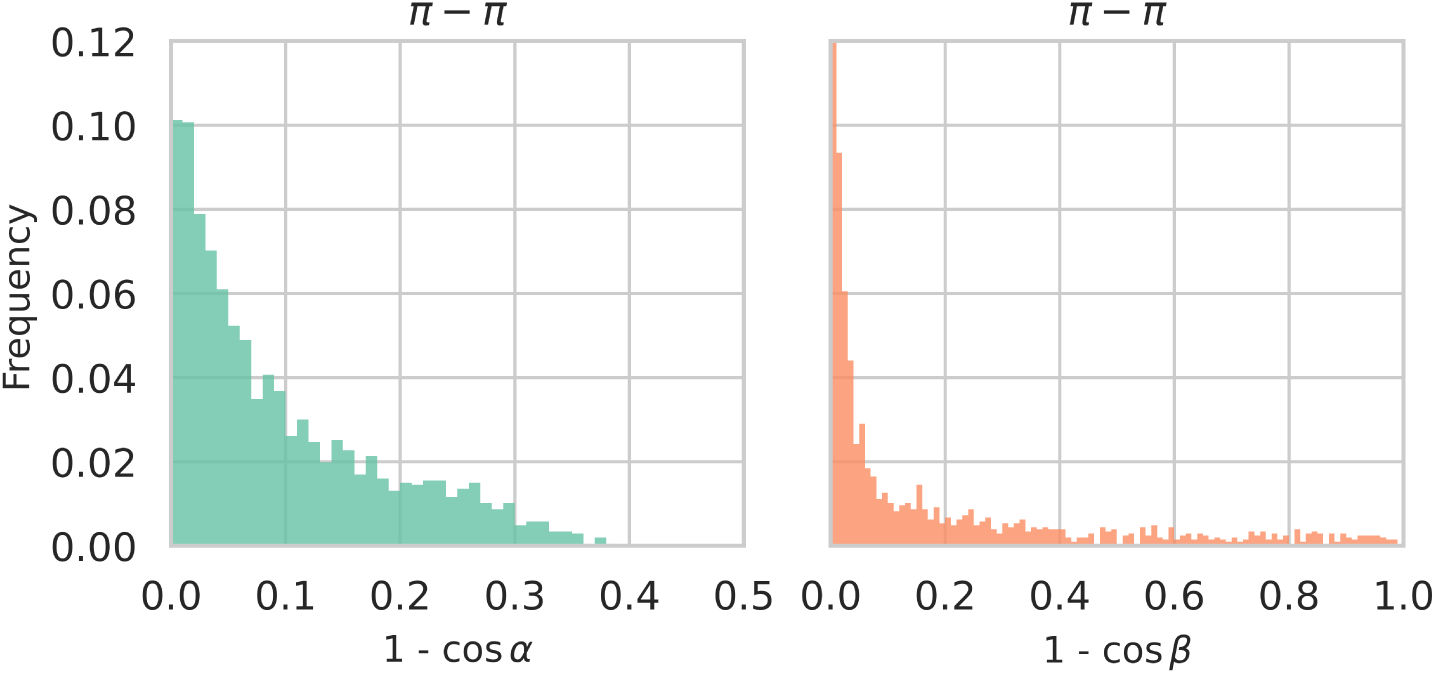
Distributions of (1*−*cos *α*) and (1*−*cos *β*) values of *π*-*π* interactions in the *D*_fibril_ set.

### 3.5 PInteract: fast, scalable, and user-friendly

PInteract is extremely easy to install and use — it only requires a standard GCC compiler and can be set up simply by cloning our GitHub repository. It is implemented in C, which makes it exceptionally fast and well-suited for large-scale analyses involving millions of structures. PInteract scales efficiently with the size of the dataset: a few thousands of structures can be processed in less than a minute on a standard laptop, and 100,000 structures require only 6 minutes (for details, see Supplementary Section S5). This level of efficiency makes PInteract particularly valuable for high-throughput structural screening tasks, as we have also highlighted in the previous sections.

## 4 Conclusion

With the recent breakthroughs in deep learning-based methods for biomolecular structure prediction, an unprecedented number of structures have been generated. We now have access to over a billion predicted protein structures, covering virtually the entire tree of life. This explosion of structural data calls for computational tools capable of efficiently analyzing these structures and extracting meaningful biological insights.

Here, we introduced PInteract, a fast computational tool designed to identify individual *π* interactions and clusters or chains of *π* interactions in proteins and in protein-protein, protein-RNA, protein-DNA, and proteinligand complexes. Moreover, PInteract is the only automatic program that identifies stair motifs at protein-DNA and protein-RNA interfaces, which are recurrent motifs combining specific H-bonds and *π* interactions. We illustrated PInteract’s capabilities across several case studies, demonstrating its effectiveness in detecting *π* interactions in proteins as well as in protein complexes. We also showed that PInteract can perform large-scale computations on hundreds of thousands of protein structures within minutes, enabling the analysis of the geometric properties of *π*-involving interactions across large structural databases. Finally, we demonstrated how PInteract can be used to gain meaningful insights into the role of *π* interactions in protein solubility and aggregation.

In summary, we are confident that PInteract, with its ease of use and high computational efficiency, will serve as a valuable resource for the scientific community, enabling the systematic study of *π*-involving interactions and shedding light on the wide spectrum of molecular mechanisms in which these interactions play a pivotal role, from protein folding and stability to biomolecular recognition and aggregation.

## Author contributions

Conceptualization, F.P. and M.R.; methodology, M.R.; software, M.R.; validation, D.L., F.P. and M.R.; data curation, D.L.; writing—original draft preparation, D.L.; writing—review and editing, F.P. and M.R. All authors have read and agreed to the published version of the manuscript.

## References

[1] Laura M Salonen, Manuel Ellermann, and Fraņcois Diederich. Aromatic rings in chemical and biological recognition: energetics and structures. Angewandte Chemie International Edition, 50(21):4808–4842, 2011.

[2] Rivka Calinsky and Yaakov Levy. Aromatic residues in proteins: Re-evaluating the geometry and energetics of *π*–*π*, cation-*π*, and ch-*π* interactions. The Journal of Physical Chemistry B, 128(36):8687–8700, 2024.

[3] SK Burley and Gregory A Petsko. Aromatic-aromatic interaction: a mechanism of protein structure stabilization. Science, 229(4708):23–28, 1985.

[4] Dennis A Dougherty. Cation-*π* interactions in chemistry and biology: a new view of benzene, phe, tyr, and trp. Science, 271(5246):163–168, 1996.

[5] Reńe Wintjens, Jacky Líevin, Marianne Rooman, and Eric Buisine. Contribution of cation-*π* interactions to the stability of protein-dna complexes. Journal of molecular biology, 302(2):393–408, 2000.

[6] Dennis A Dougherty. The cation-*π* interaction in chemistry and biology. Chemical Reviews, 2025.

[7] Yumi N Imai, Yoshihisa Inoue, Isao Nakanishi, and Kazuo Kitaura. Amide–*π* interactions between formamide and benzene. Journal of Computational Chemistry, 30(14):2267–2276, 2009.

[8] Emilie Caüet, Marianne Rooman, Reńe Wintjens, Jacques Líevin, and Christophe Biot. Histidine-aromatic interactions in proteins and proteinligand complexes: quantum chemical study of x-ray and model structures. Journal of chemical theory and computation, 1(3):472–483, 2005.

[9] Nobuhisa Shimba, Zach Serber, Richard Ledwidge, Susan M Miller, Charles S Craik, and Volker Dötsch. Quantitative identification of the protonation state of histidines in vitro and in vivo. Biochemistry, 42(30):9227–9234, 2003.

[10] Rivka Calinsky and Yaakov Levy. Histidine in proteins: ph-dependent interplay between *π*–*π*, cation–*π*, and ch–*π* interactions. Journal of Chemical Theory and Computation, 20(15):6930–6945, 2024.

[11] KSC Reid, PF Lindley, and JM Thornton. Sulphur-aromatic interactions in proteins. FEBS letters, 190(2):209–213, 1985.

[12] William B Motherwell, Rafael B Moreno, Ilias Pavlakos, Josephine RT Arendorf, Tanzeel Arif, Graham J Tizzard, Simon J Coles, and Abil E Aliev. Noncovalent interactions of *π* systems with sulfur: The atomic chameleon of molecular recognition. Angewandte Chemie, 130(5):1207–1212, 2018.

[13] Luka Nunar and Abil E Aliev. Quantification of the strength of *π*-noncovalent interactions in molecular balances using density functional methods. Chemistry-Methods, 3(1):e202200044, 2023.

[14] Kristina N-M Daeffler, Henry A Lester, and Dennis A Dougherty. Functionally important aromatic–aromatic and sulfur-*π* interactions in the d2 dopamine receptor. Journal of the American Chemical Society, 134(36):14890–14896, 2012.

[15] Esteban Lanzarotti, Rolf R Biekofsky, Daŕıo A Estrin, Marcelo A Marti, and Adrían G Turjanski. Aromatic–aromatic interactions in proteins: beyond the dimer. Journal of chemical information and modeling, 51(7):1623–1633, 2011.

[16] Marianne Rooman, Jacky Líevin, Eric Buisine, and Reńe Wintjens. Cation–*π*/h-bond stair motifs at protein–dna interfaces. Journal of molecular biology, 319(1):67–76, 2002.

[17] Christophe Biot, Reńe Wintjens, and Marianne Rooman. Stair motifs at proteindna interfaces: nonadditivity of h-bond, stacking, and cation-*π* interactions. Journal of the American Chemical Society, 126(20):6220–6221, 2004.

[18] Kiran Kumar, Shin M. Woo, Thomas Siu, Wilian A. Cortopassi, Fernanda Duarte, and Robert S. Paton. Cation–*π* interactions in protein–ligand binding: theory and data-mining reveal different roles for lysine and arginine. Chemical Science, 9(10):2655–2665, 2018.

[19] Nandan Kumar, Anamika Singh Gaur, and G Narahari Sastry. A perspective on the nature of cation-*π* interactions. Journal of Chemical Sciences, 133(4):97, December 2021.

[20] Dhawal Shah, Jianguo Li, Abdul Rajjak Shaikh, and Raj Rajagopalan. Arginine–aromatic interactions and their effects on arginine-induced solubilization of aromatic solutes and suppression of protein aggregation. Biotechnology Progress, 28(1):223–231, January 2012.

[21] R. J. Zauhar, C. L. Colbert, R. S. Morgan, and W. J. Welsh. Evidence for a strong sulfur-aromatic interaction derived from crystallographic data. Biopolymers, 53(3):233–248, March 2000.

[22] Georg Krainer, Timothy J. Welsh, Jerelle A. Joseph, Jorge R. Espinosa, Sina Wittmann, Ella De Csilĺery, Akshay Sridhar, Zenon Toprakcioglu, Giedre Gudišskytė, Magdalena A. Czekalska, William E. Arter, Jordina Guilĺen-Boixet, Titus M. Franzmann, Seema Qamar, Peter St George-Hyslop, Anthony A. Hyman, Rosana Collepardo-Guevara, Simon Alberti, and Tuomas P. J. Knowles. Reentrant liquid condensate phase of proteins is stabilized by hydrophobic and non-ionic interactions. Nature Communications, 12(1):1085, February 2021.

[23] Hongxia Zhao, Ling Tang, Yu Fang, Chao Liu, Wenlong Ding, Shunping Zang, Yulin Chen, Wenyuan Xu, Ying Yuan, Dong Fang, and Shixian Lin. Manipulating Cation*π* Interactions of Reader Proteins in Living Cells with Genetic Code Expansion. Journal of the American Chemical Society, 145(30):16406–16416, August 2023.

[24] Christopher C. Valley, Alessandro Cembran, Jason D. Perlmutter, Andrew K. Lewis, Nicholas P. Labello, Jiali Gao, and Jonathan N. Sachs. The Methionine-aromatic Motif Plays a Unique Role in Stabilizing Protein Structure. Journal of Biological Chemistry, 287(42):34979–34991, October 2012.

[25] Y. Bhargav Kumar, Nandan Kumar, Lijo John, Hridoy Jyoti Mahanta, S. Vaikundamani, Selvaraman Nagamani, G. Madhavi Sastry, and G. Narahari Sastry. Analyzing the cation-aromatic interactions in proteins: Cation-aromatic database V2.0. Proteins: Structure, Function, and Bioinformatics, 92(2):179–191, February 2024.

[26] Mukesh Chourasia, G. Madhavi Sastry, and G. Narahari Sastry. Aromatic–Aromatic Interactions Database, A2ID: An analysis of aromatic π-networks in proteins. International Journal of Biological Macromolecules, 48(4):540–552, May 2011.

[27] K. G. Tina, R. Bhadra, and N. Srinivasan. PIC: Protein Interactions Calculator. Nucleic Acids Research, 35(Web Server):W473–W476, May 2007.

[28] Hervé Minoux and Christophe Chipot. Cation-*π* Interactions in Proteins: Can Simple Models Provide an Accurate Description? Journal of the American Chemical Society, 121(44):10366–10372, November 1999.

[29] Justin P. Gallivan and Dennis A. Dougherty. A Computational Study of Cation-*π* Interactions vs Salt Bridges in Aqueous Media: Implications for Protein Engineering. Journal of the American Chemical Society, 122(5):870–874, February 2000.

[30] R. J. Zauhar, C. L. Colbert, R. S. Morgan, and W. J. Welsh. Evidence for a strong sulfur-aromatic interaction derived from crystallographic data. Biopolymers, 53(3):233–248, March 2000. Publisher: Wiley.

[31] Guoli Wang and Roland L Dunbrack Jr. Pisces: a protein sequence culling server. Bioinformatics, 19(12):1589–1591, 2003.

[32] Ragul Gowthaman and Brian G Pierce. Tcr3d: The t cell receptor structural repertoire database. Bioinformatics, 35(24):5323–5325, 2019.

[33] Brian G Pierce and Zhiping Weng. A flexible docking approach for prediction of t cell receptor–peptide–mhc complexes. Protein Science, 22(1):35–46, 2013.

[34] James Dunbar, Konrad Krawczyk, Jinwoo Leem, Terry Baker, Angelika Fuchs, Guy Georges, Jiye Shi, and Charlotte M Deane. Sabdab: the structural antibody database. Nucleic acids research, 42(D1):D1140–D1146, 2014.

[35] Constantin Schneider, Matthew IJ Raybould, and Charlotte M Deane. Sabdab in the age of biotherapeutics: updates including sabdab-nano, the nanobody structure tracker. Nucleic acids research, 50(D1):D1368–D1372, 2022.

[36] Matthew IJ Raybould, Claire Marks, Alan P Lewis, Jiye Shi, Alexander Bujotzek, Bruck Taddese, and Charlotte M Deane. Thera-sabdab: the therapeutic structural antibody database. Nucleic acids research, 48(D1):D383–D388, 2020.

[37] Helen M Berman, John Westbrook, Zukang Feng, Gary Gilliland, Talapady N Bhat, Helge Weissig, Ilya N Shindyalov, and Philip E Bourne. The protein data bank. Nucleic acids research, 28(1):235–242, 2000.

[38] Georgios A Dalkas, Fabian Teheux, Jean Marc Kwasigroch, and Marianne Rooman. Cation–*π*, amino–*π*, *π*–*π*, and h-bond interactions stabilize antigen–antibody interfaces. Proteins: Structure, Function, and Bioinformatics, 82(9):1734–1746, 2014.

[39] Hiroki Akiba and Kouhei Tsumoto. Thermodynamics of antibody–antigen interaction revealed by mutation analysis of antibody variable regions. The Journal of Biochemistry, 158(1):1–13, July 2015.

[40] Ian K McDonald and Janet M Thornton. Satisfying hydrogen bonding potential in proteins. Journal of molecular biology, 238(5):777–793, 1994.

[41] Di Wu, Jing Sun, Tianlei Xu, Shuning Wang, Guoqing Li, Yixue Li, and Zhiwei Cao. Stacking and energetic contribution of aromatic islands at the binding interface of antibody proteins. In Immunome research, volume 6, pages 1–9. Springer, 2010.

[42] Mihaly Varadi, Damian Bertoni, Paulyna Magana, Urmila Paramval, Ivanna Pidruchna, Malarvizhi Radhakrishnan, Maxim Tsenkov, Sreenath Nair, Milot Mirdita, Jingi Yeo, et al. Alphafold protein structure database in 2024: providing structure coverage for over 214 million protein sequences. Nucleic acids research, 52(D1):D368–D375, 2024.

[43] John Jumper, Richard Evans, Alexander Pritzel, Tim Green, Michael Figurnov, Olaf Ronneberger, Kathryn Tunyasuvunakool, Russ Bates, Augustin Žídek, Anna Potapenko, et al. Highly accurate protein structure prediction with alphafold. nature, 596(7873):583–589, 2021.

[44] Qingzhen Hou, Raphäel Bourgeas, Fabrizio Pucci, and Marianne Rooman. Computational analysis of the amino acid interactions that promote or decrease protein solubility. Scientific reports, 8(1):14661, 2018.

[45] Robert McCoy Vernon, Paul Andrew Chong, Brian Tsang, Tae Hun Kim, Alaji Bah, Patrick Farber, Hong Lin, and Julie Deborah Forman-Kay. Pi-Pi contacts are an overlooked protein feature relevant to phase separation. eLife, 7:e31486, February 2018.

[46] Jim Warwicker, Spyros Charonis, and Robin A. Curtis. Lysine and Arginine Content of Proteins: Computational Analysis Suggests a New Tool for Solubility Design. Molecular Pharmaceutics, 11(1):294–303, January 2014. Publisher: American Chemical Society (ACS).

[47] Ehud Gazit. A possible role for *π*-stacking in the self-assembly of amyloid fibrils. The FASEB Journal, 16(1):77–83, 2002.

[48] Qingzhen Hou, Jean Marc Kwasigroch, Marianne Rooman, and Fabrizio Pucci. Solart: a structure-based method to predict protein solubility and aggregation. Bioinformatics, 36(5):1445–1452, 2020.

[49] Tatsuya Niwa, Bei-Wen Ying, Katsuyo Saito, WenZhen Jin, Shoji Takada, Takuya Ueda, and Hideki Taguchi. Bimodal protein solubility distribution revealed by an aggregation analysis of the entire ensemble of escherichia coli proteins. Proceedings of the National Academy of Sciences, 106(11):4201–4206, 2009.

[50] Eri Uemura, Tatsuya Niwa, Shintaro Minami, Kazuhiro Takemoto, Satoshi Fukuchi, Kodai Machida, Hiroaki Imataka, Takuya Ueda, Motonori Ota, and Hideki Taguchi. Large-scale aggregation analysis of eukaryotic proteins reveals an involvement of intrinsically disordered regions in protein folding. Scientific reports, 8(1):678, 2018.

[51] Jim Warwicker, Spyros Charonis, and Robin A Curtis. Lysine and arginine content of proteins: computational analysis suggests a new tool for solubility design. Molecular pharmaceutics, 11(1):294–303, 2014.

[52] Ivana M Stankovíc, Shuqiang Niu, Michael B Hall, and Snězana D Zaríc. Role of aromatic amino acids in amyloid self-assembly. International journal of biological macromolecules, 156:949–959, 2020.

[53] Rob van der Kant, Nikolaos Louros, Joost Schymkowitz, and Frederic Rousseau. Thermodynamic analysis of amyloid fibril structures reveals a common framework for stability in amyloid polymorphs. Structure, 30(8):1178–1189, 2022.

