## Supplementary material for "PInteract: Detecting Aromatic-Involving Motifs in Proteins and Protein-Nucleic Acid Complexes": Supplemtary Material

Computational Biology and Bioinformatics, Université Libre de Bruxelles, Belgium
and
Interuniversity Institute of Bioinformatics in Brussels, Belgium

### Section S1. Relative frequencies of $\pi$-interactions in different datasets

Table S1: P-values associated to the t-tests of the average relative frequencies of $\pi$-interactions between the four datasets $\mathcal{D}_{\text{monomer}}$, $\mathcal{D}_{\text{dimer}}$, $\mathcal{D}_{\text{TCRpMHC}}$ and $\mathcal{D}_{\text{AbAg}}$. These frequencies are computed as the number of interactions identified by PInteract divided by the total number of interactions in the protein. For the frequencies and computation details, see Table 1 of the main manuscript.

| | Cation-$\pi$ | Amino-$\pi$ | His-$\pi$ | Sulfur-$\pi$ | $\pi$-$\pi$ |
|---|---|---|---|---|---|
| $\mathcal{D}_{\text{monomer}}$-$\mathcal{D}_{\text{dimer}}$ | $3.63\times10^{-7}$ | 0.12 | $1.33\times10^{-7}$ | 0.91 | 0.62 |
| $\mathcal{D}_{\text{monomer}}$-$\mathcal{D}_{\text{TCRpMHC}}$ | 0.25 | 0.03 | 0.40 | 0.08 | 0.002 |
| $\mathcal{D}_{\text{monomer}}$-$\mathcal{D}_{\text{AbAg}}$ | $3.78\times10^{-16}$ | $1.51\times10^{-12}$ | $3.33\times10^{-6}$ | $9.33\times10^{-5}$ | 0.02 |
| $\mathcal{D}_{\text{dimer}}$-$\mathcal{D}_{\text{TCRpMHC}}$ | 0.62 | 0.05 | 0.08 | 0.08 | 0.003 |
| $\mathcal{D}_{\text{dimer}}$-$\mathcal{D}_{\text{AbAg}}$ | $3.20\times10^{-9}$ | $7.57\times10^{-9}$ | $2.06\times10^{-10}$ | $2.71\times10^{-4}$ | 0.03 |
| $\mathcal{D}_{\text{TCRpMHC}}$-$\mathcal{D}_{\text{AbAg}}$ | $1.21\times10^{-5}$ | 0.77 | 0.31 | 0.95 | 0.12 |

### Section S2. Distributions of $\alpha$ and $\beta$ angles characterizing $\pi$ interaction geometry in experimental structures

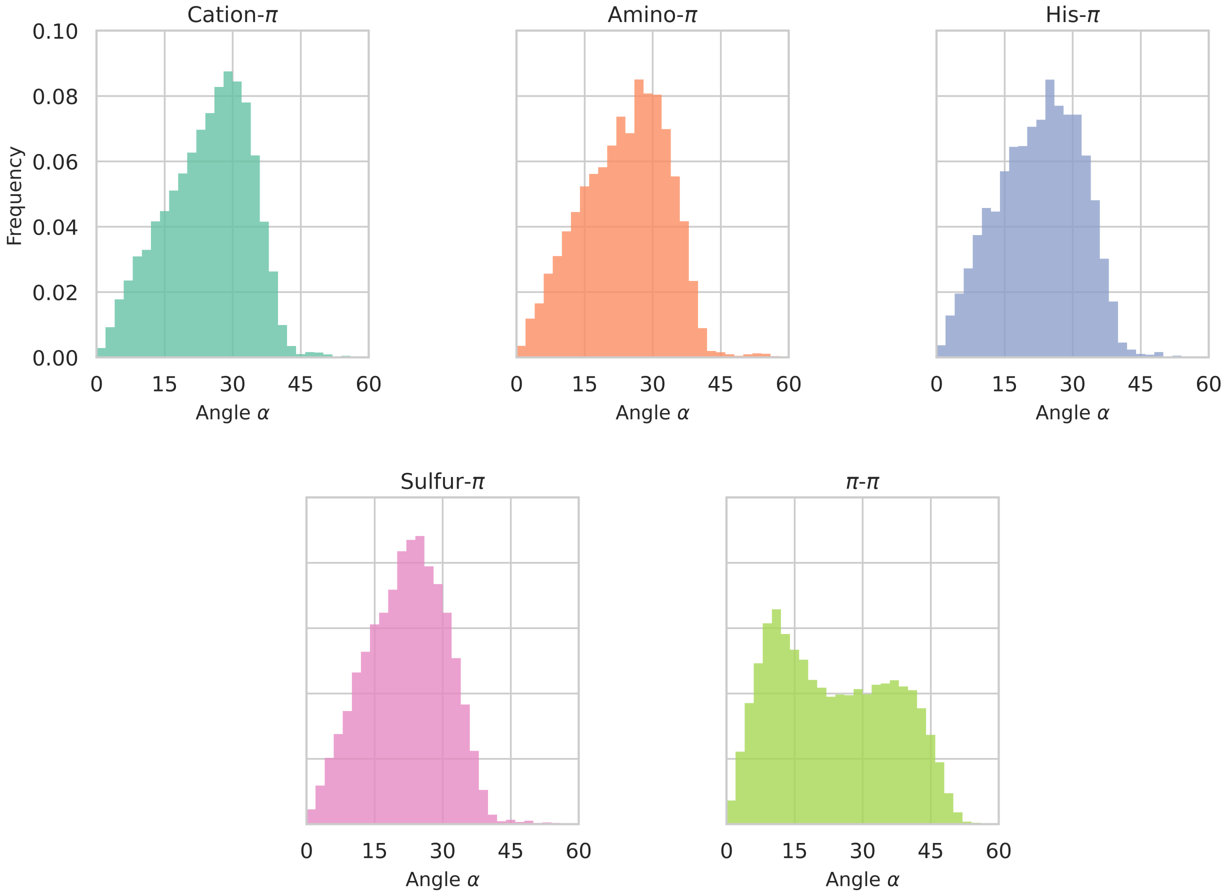


Figure S1: Distribution of $\alpha$ angle values measuring the location of the functional group of interacting partner 2 above (or below) the plane of the aromatic ring of partner 1, for different types of $\pi$ interactions detected in the $\mathcal{D}_{\text{monomer}}$ dataset. The smaller the value of $\alpha$, the more directly the functional group of partner 2 is positioned above (or below) the center of the aromatic ring. See Figure 5 of the main manuscript for details and the distribution of the lateral displacement with respect to the aromatic rings' center, $(1 - \cos\alpha)$.

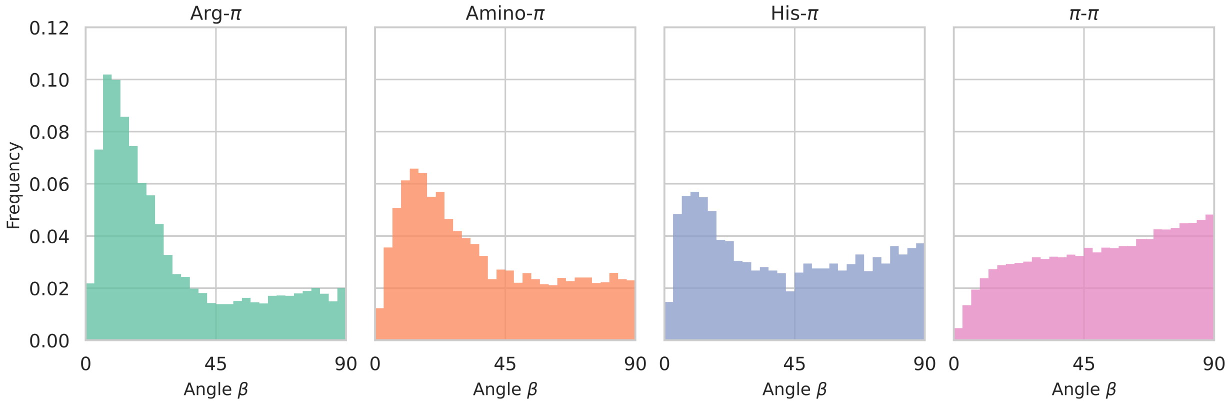


Figure S2: Distribution of $\beta$ angle values measuring the degree of parallelism between the interacting planar functional groups, for different types of $\pi$ interactions detected in the $\mathcal{D}_{\text{monomer}}$ dataset. Parallel or stacked conformations correspond to $\beta = 0°$; perpendicular or T-shaped conformations, to $\beta = 90°$. See Figure 6 of the main manuscript for details and the distribution of the displacement values $(1 - \cos\beta)$.

### Section S3. Distributions of $\alpha$ and $\beta$ angles characterizing $\pi$ interaction geometry in AlphaFold-predicted structures

To investigate whether the angular preferences of $\pi$ interactions are also retained in computationally predicted structures, we applied PInteract to representative proteins from the AlphaFold Protein Structure Database (AlphaFoldDB) [1]. We assembled a dataset, denoted $\mathcal{D}_{\mathrm{AF}}$, comprising representative subsets of proteomes from four model organisms in AlphaFoldDB: *Arabidopsis thaliana*, *Escherichia coli*, *Homo sapiens*, and *Saccharomyces cerevisiae*. For each protein in $\mathcal{D}_{\mathrm{AF}}$, we run PInteract and extracted all $\pi$-involving interactions and the corresponding $\alpha$ and $\beta$ angles as described in the main manuscript. The results are shown in Figure S3 and S4.

The distributions of $\alpha$ angles shown in Figure S3 are highly similar to those observed in experimentally determined monomer structures (see Figure 5 of the main manuscript), indicating that AlphaFold [2] tends to correctly predict $\pi$ interaction geometries in which the interaction partner is located above (or below) the aromatic ring.

The $\beta$ angle distributions depicted in Figure S4 also largely resemble those of experimental protein structures (see Figure 6 of the main manuscript), except for a subtle but systematic reduction of the frequency of stacked conformations for Arg-$\pi$, amino-$\pi$, and His-$\pi$ interactions. In $\mathcal{D}_{\mathrm{AF}}$, stacked conformations are less representated than in experimental protein striuctures, whereas T-shaped confirmations are slightly overrepresented (Figure S4). This suggests that AlphaFold underestimates the energetic stabilization associated with stacked conformations in certain $\pi$-interactions.

Despite this minor difference, the general similarity between the angular distributions of $\pi$ interactions in AlphaFoldDB and those observed in experimental protein structures supports the geometric realism of AlphaFold-predicted $\pi$ interactions and further demonstrates the applicability of PInteract to computationally predicted structures.

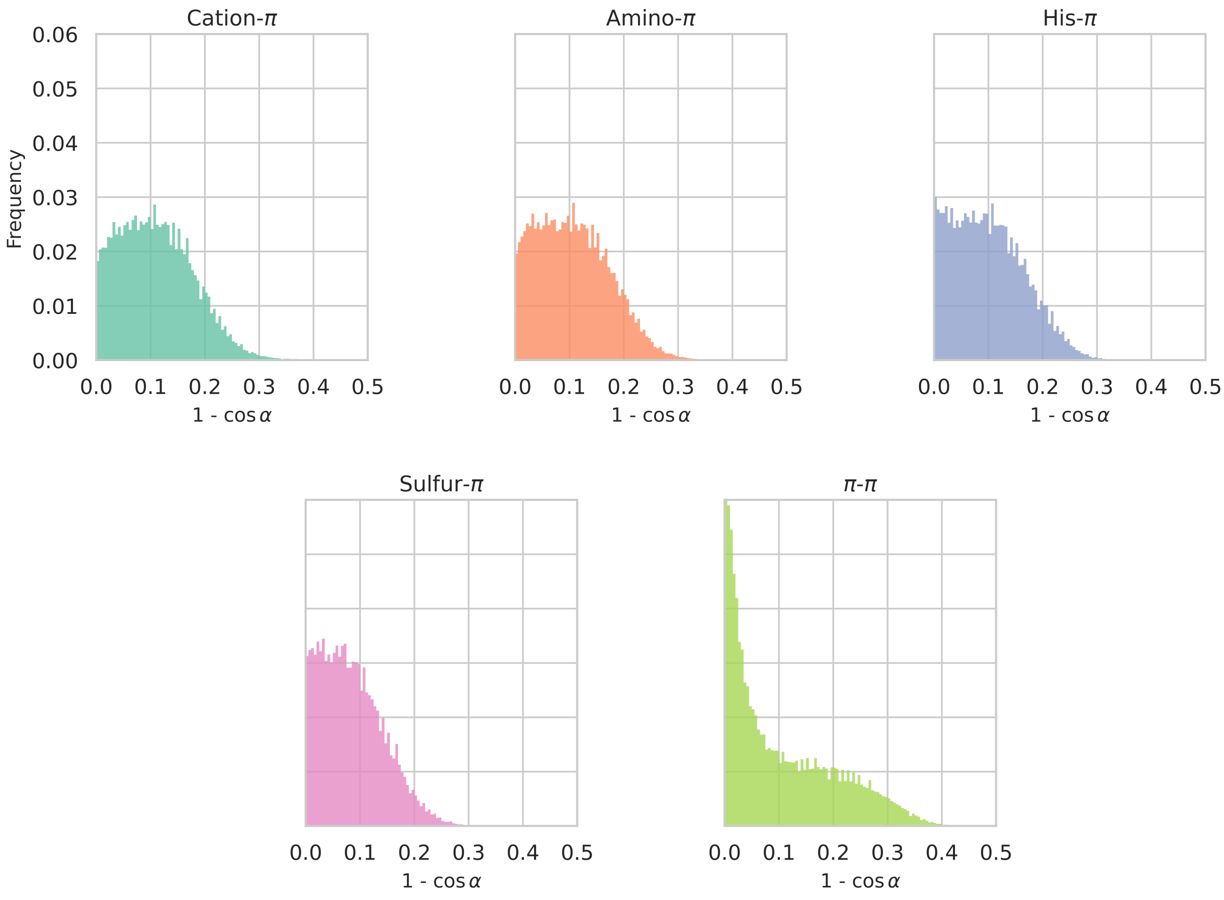


Figure S3: Distribution of the $1 - \cos\angle$ values of the lateral displacement with respect to the aromatic rings' center, $(1 - \cos\alpha)$, in which the $\alpha$ angle measures the location of the functional group of interacting partner 2 above (or below) the plane of the aromatic ring of partner 1, for different types of $\pi$ interactions detected in the $\mathcal{D}_{\mathrm{AF}}$ dataset. The smaller the value of $(1 - \cos\alpha)$, the more directly the functional group of partner 2 is positioned above (or below) the center of the aromatic ring. This Figure must be compared with Figure 5 in the main manuscript that show the equivalent distributions in experimental structures.

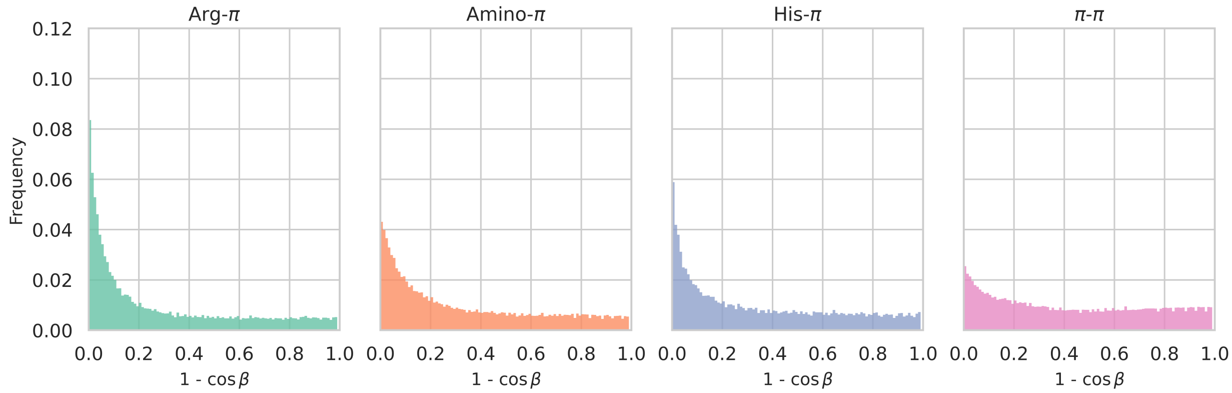


Figure S4: Distribution of $1 - \cos\beta$ values, in which the $\beta$ angle measures the degree of parallelism between the interacting planar functional groups, for different types of $\pi$ interactions detected in the $\mathcal{D}_{\mathrm{AF}}$ dataset. Parallel or stacked conformations correspond to $\beta = 0°$ and $1 - \cos\beta = 0$; perpendicular or T-shaped conformations, to $\beta = 90°$ and $1 - \cos\beta = 1$. This Figure must be compared with Figure 6 in the main manuscript that show the equivalent distributions in experimental structures.

### Section S4. Example of $\pi$-$\pi$ chains in amyloid fibrils

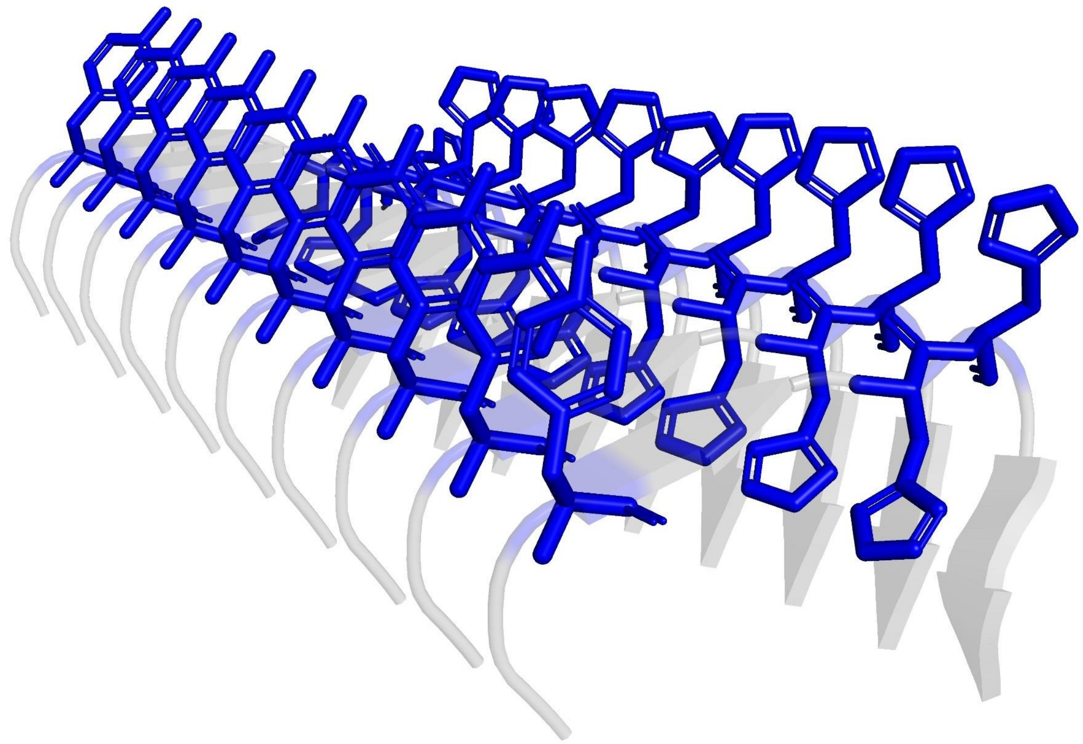

Figure S5: $\pi$-$\pi$ Chain in an A$\beta$ amyloid fibril purified from Alzheimer's brain tissue (PDB id: 6SHS) [3].

### Section S5. Running efficiency of PInteract

Table S2: Running time of PInteract on datasets of different size.

| Dataset | Number of samples | Time (seconds) |
|---|---|---|
| *Arabidopsis thaliana* | 27,434 | 90.1 |
| *Escherichia coli* | 4,363 | 9.5 |
| *Homo sapiens* | 23,391 | 129.4 |
| *Saccharomyces cerevisiae* | 6,039 | 24.0 |

### References


[1] Mihaly Varadi, Damian Bertoni, Paulyna Magana, Urmila Paramval, Ivanna Pidruchna, Malarvizhi Radhakrishnan, Maxim Tsenkov, Sreenath Nair, Milot Mirdita, Jingi Yeo, et al. Alphafold protein structure database in 2024: providing structure coverage for over 214 million protein sequences. *Nucleic acids research*, 52(D1):D368–D375, 2024.

[2] John Jumper, Richard Evans, Alexander Pritzel, Tim Green, Michael Figurnov, Olaf Ronneberger, Kathryn Tunyasuvunakool, Russ Bates, Augustin Žídek, Anna Potapenko, et al. Highly accurate protein structure prediction with alphafold. *nature*, 596(7873):583–589, 2021.

[3] Marius Kollmer, William Close, Leonie Funk, Jay Rasmussen, Aref Bsoul, Angelika Schierhorn, Matthias Schmidt, Christina J Sigurdson, Mathias Jucker, and Marcus Fändrich. Cryo-em structure and polymorphism of a$\beta$ amyloid fibrils purified from alzheimer's brain tissue. *Nature communications*, 10(1):4760, 2019.
